# Trophic complexity alters the diversity-multifunctionality relationship in experimental grassland mesocosms

**DOI:** 10.1101/525881

**Authors:** Krishna Anujan, Sebastian A. Heilpern, Case M. Prager, Brian C. Weeks, Shahid Naeem

## Abstract

Diversity within trophic levels influences the number of ecosystem functions maintained simultaneously by a community, or multifunctionality. Depending on threshold cutoffs applied to measuring these functions, the diversity-multifunctionality relationship changes from positive until intermediate thresholds to negative effect at high function thresholds. Although the presence of multiple trophic levels or trophic complexity affects levels of functions, its effect on the diversity-multifunctionality relationship has not been experimentally tested. We simultaneously manipulated plant diversity and trophic complexity in a multifactorial tall-grass prairie mesocosm experiment at Cedar Creek, Minnesota, USA and measured multiple ecosystem functions. Trophic complexity altered the diversity-multifunctionality relationship in two key ways: it lowered the maximum strength of the diversity-multifunctionality effect and it resulted in a switch from positive to negative relationship between increasing diversity and multifunctionality at lower function thresholds. Our findings suggest that global declines in trophic complexity will exacerbate the reduction in ecosystem multifunctionality as a result of widespread declines in biodiversity.

## Introduction

Increasing biodiversity within a trophic level, both along experimental and naturally occurring gradients, is associated with a positive and saturating increase in the magnitude of ecosystem functions when considered individually and often a monotonic increase in the number of functions maintained simultaneously^1–5^. Empirical support for this diversity-multifunctionality relationship, across taxa and habitats, suggests that higher levels of biodiversity may be necessary to maintain ecosystem functioning than previously assumed based on single-function studies^2,6,7^.

Multifunctionality of ecosystems is sensitive to several factors, prominently: (1) the levels of biodiversity within trophic levels^4,7^ and (2) a desired “threshold” magnitude of each ecosystem function scaled from zero to 100% (above this level, a community is considered to maintain that particular function and contribute to multifunctionality)^7–9^. When low thresholds are considered (e.g., maintining at least 10% of every function measured) diversity generally increases the number of functions maintained above that threshold; a positive diversity-multifunction relationship. But this effect of diversity on multifunctionality, or slope of this relationship between number of ecosystem functions maintained and diversity, is expected to increase with the selected threshold only until moderate threshold values, then decreases and switches to a negative diversity-multifunction relationship at high thresholds. This pattern of changes in the diversity-multifunctionality relationship is referred to as the “jack-of-all-trades” effect^8^ (Fig 1). In effect, community average values for functions are expected to be low in diverse communities compared to high functioning monocultures, allowing them to maintain most functions, but only at the average value contributed by each species.

**Figure 1.**
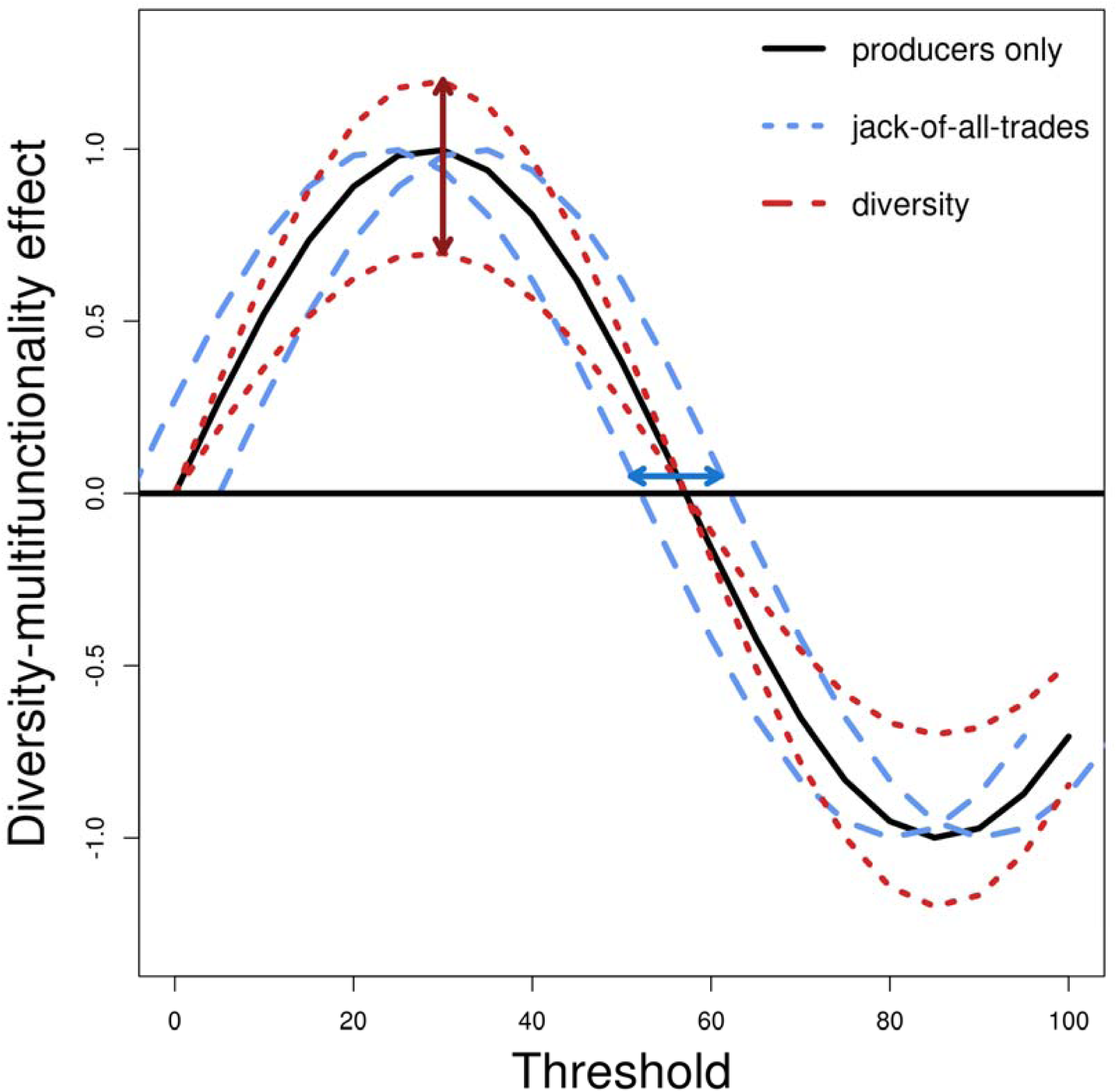
Conceptual framework illustrating hypothesised effects of trophic complexity on the diversity-multifunctionality effect (DME) curve. The DME is a measure of the slope of the relationship between diversity and multifunctionality (e.g., number of functions gained through the addition of species) whose value is dependent on the threshold (i.e., percent ecosystem function obtained) used in estimating multifunctionality. Positive effects of diversity correspond to positive DME values, or a curve above zero and vice versa. The continuous black curve represents a hypothetical relationship between selected threshold value and DME for a plant community in the absence of trophic complexity (i.e., the plant only curve). The red and blue lines represent possible deviations from the plant-only curve with the addition of trophic complexity. The red curves represent trophic-induced changes to diversity effects on single ecosystem funcions, which alter the flatness of the DME curve. The blue curves represent trophic-induced changes to correlations between traits or altered “jack-of-all-trades” effect, which shifts the horizontal location of the DME switch from positive to negative.

Most diversity-function studies and tests of this pattern have focussed on a single trophic level, most often the plant or producer community, but changes in diversity of non-producer trophic levels can also have significant impacts on several ecosystem functions^10–16^. Plant diversity can affect diversity and community composition of both aboveground and litter fauna which by removing biomass and through decomposition, can then affect ecosystem functioning. However, non-producer trophic levels can either enhance the effect of plant diversity on an ecosystem function^17^ (e.g. nutrient uptake) or reduce it^18^ (e.g. primary productivity). The number of trophic levels, or trophic complexity, can thus potentially impact the diversity-multifunctionality relationship, but this effect is poorly understood. Current studies do not allow for a direct comparison of ecosystems that vary in trophic complexity because such explorations would require simultaneously manipulating plant diversity as well as the number of non-producer trophic groups.

Here, we explore the effects of diversity and trophic complexity on ecosystem multifunctionality using an experimental approach. In the absence of any effect by non-prodcuer trophic groups, we hypothesise that plant diversity would increase multifunctionality until moderate thresholds and decrease multifunctionality at high thresholds, resulting in a jack-of-all-trades relationship. We hypothesise that trophic complexity could have impacts on multifunctionality at two different levels; (1) altered diversity-function relationships for single functions – the number and identity of non-producer trophic levels affects each ecosystem function distinctly, either strengthening or dampening the effect of plant diversity which aggregates to strengthened or dampened multifunctionality effects and (2) non-producer trophic level affect plant traits at the species level altering correlations between measured functions. If trait trade-offs are reduced in the presence of multiple trophic levels, communities would maintain multiple functions for higher than moderate thresholds and vice versa. This shift the threshold at which the diversity-multifunctionality effect switches from positive to negative (Heilpern et al. in review). Each of these effects can be measured on the jack-off-all-trades curve, or the relationship between the strength (slope) of the diversity-multifunction effect (DME) and the selected threshold for measuring multifunctionality. Based on the hypotheses, we expected two corresponding effects of trophic complexity to this curve: (1) a change in the magnitude of the DME across thresholds, resulting in either a taller or flatter curve (2) a change in the threshold at which DME shifts from positive to negative effect measured along the horizontal location on the x-axis. Figure 1 provides a conceptual framework for these predicted outcomes. We note that these predicted responses to trophic complexity are not mutually exclusive; the curve flatness, shift location, both, or neither may respond to differences in trophic complexity.

To test this framework, we simultaneously manipulated tall-grass prairie plant diversity and trophic complexity in 94 tall-grass prairie mesocosms at Cedar Creek, Minnesota, USA. In these mesocosms, we varied plant diversity from 1 to 16 species, following a standard, stratified log_2_ randomised design. We simultaneously varied trophic complexity following a factorial design resulting in 4 trophic treatments; above-ground insect-dominated mesofaunal communities only (INS), below-ground litter mesofaunal communities only (LIT), both aboveground and litter mesofauna (BOTH) or no non-producer trophic levels (NONE). For comparison, we pooled all the data (POOLED).

We measured four ecosystem functions; aboveground biomass, belowground biomass, soil water retention and biomass recovery after harvest. We calculated the mean values of these functions across the different plant diversity and trophic complexity treatments. We standardised the values of each of the functions between 0 and 100 for the entire dataset and calculated a combined multifunctionality metric, the number of functions maintained above a given threshold for each community across this range of thresholds following the standard approach^4,8^. We then analysed the diversity-multifunctionality effect (the DME) as the slope of the linear fit of the number of functions maintained above the threshold against plant diversity. Finally, we tested whether this relationship was sensitive to trophic complexity.

## Results

For every function measured, communities on average, independent of diversity and trophic complexity treatments, had low values; most communities failed to maintain functions at high values (Fig 2). The average values of each function remained within a small range of values across different plant diversity treatments (Fig S1). Trophic treatment did not alter the curves substantially for any single function, although there were some differences at high threshold values for water retention and biomass recovery (Fig 2) and at high diversity treatments for water retention and aboveground biomass (Fig S1).

**Figure 2:**
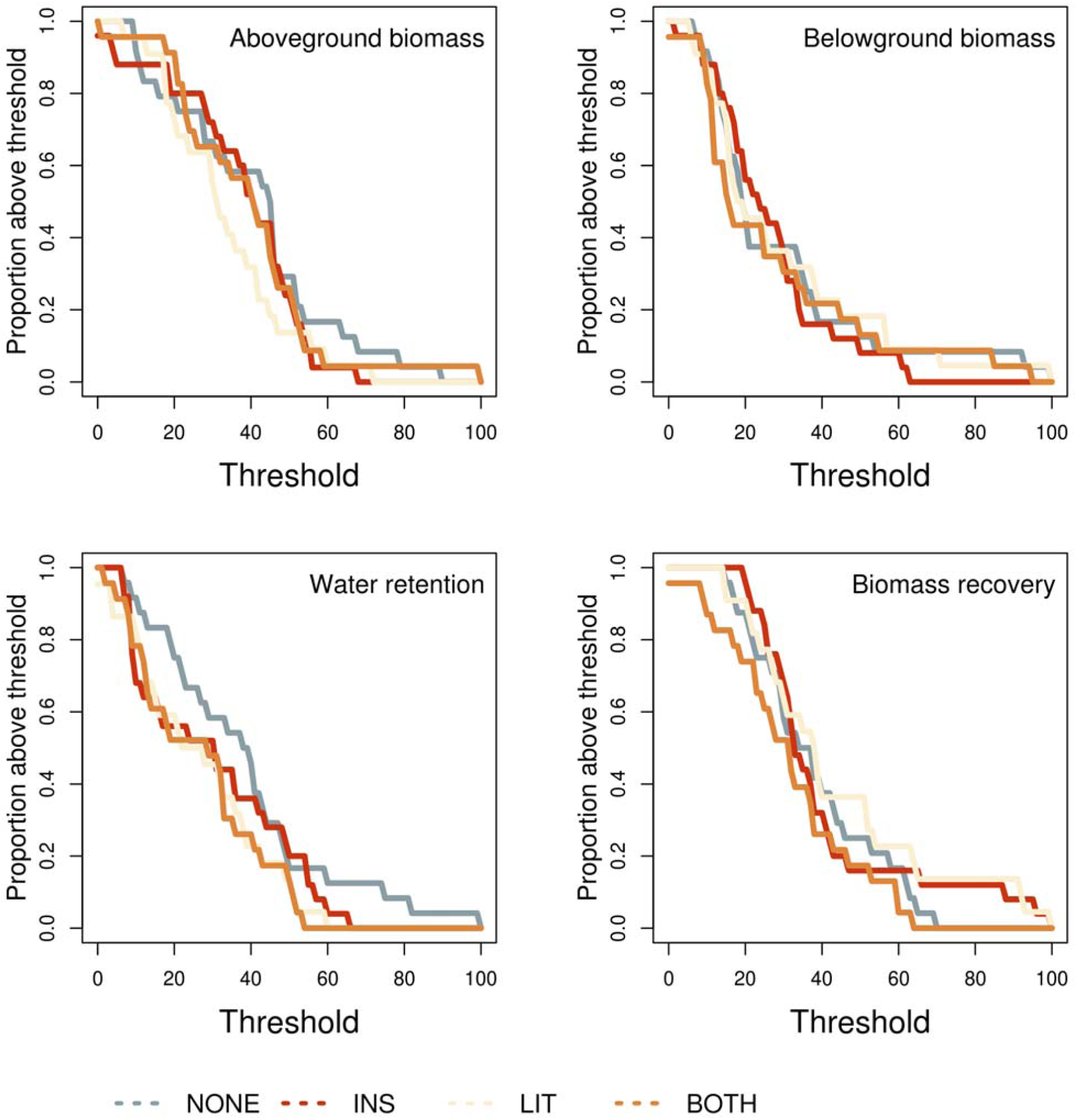
Proportion of communities maintaining single ecosystem functions along a range of percent thresholds. Panels represent the four functions – aboveground biomass, belowground biomass, water retention and biomass recovery. Coloured curves represent the four trophic treatments; Only plants (NONE, grey), plants and aboveground mesofauna (INS, red), plants and litter mesofauna (LIT, cream), plants and both aboveground and litter mesofauna (BOTH, orange).

Ecosystem multifunctionality showed the predicted pattern to changes in plant biodiversity, with positive effects at low thresholds and negative effects at high thresholds (Fig. 3). The plant diversity-multifunctionality relationship for the pooled dataset was positive for a threshold of 25% (slope=0.03, p=0.36) and 50% (slope=0.007, p=0.81) but was negative for 75% (slope=-0.14, p=0.4) and 90% thresholds (slope=-0.11, p=0.36) (Fig 3). At the 90% threshold, most plots had no functions above cutoff, making a biodiversity or trophic effect less discernible, whereas at 25% threshold, most communities mauntained functions even without the effect of plant diversity or trophic structure. The biodiversity effect on multifunctionality was most observable at moderate thresholds. Similarly, the effect of trophic complexity was significant only at 75% (−0.12, p<0.05) and marginally significant at 90% (−0.07, p=0.06) while at a 25% threshold (−0.05, p=0.59) and 50% threshold (−0.14, p=0.16), these differences were not detectable.

**Figure 3.**
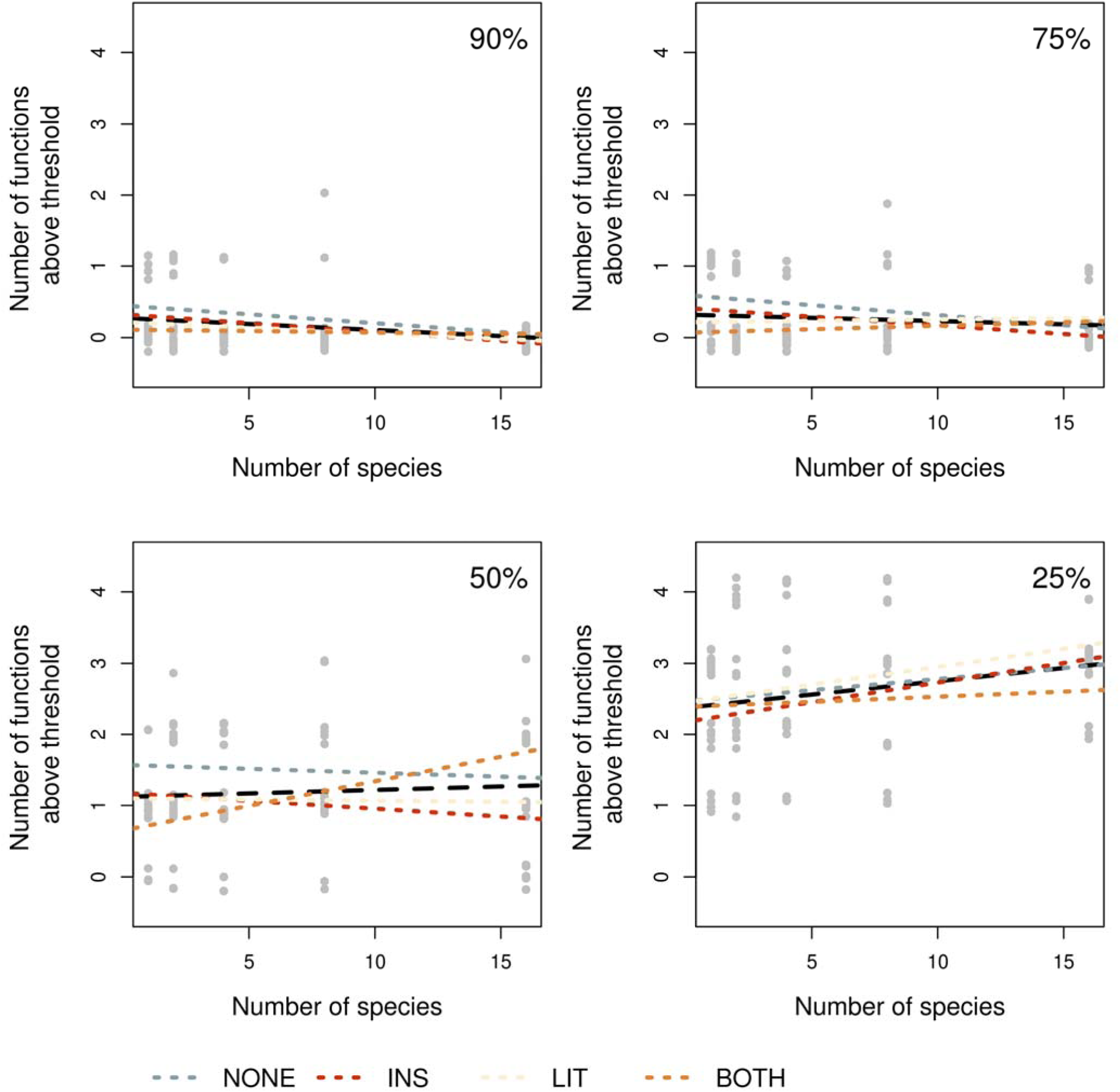
Number of functions above four different thresholds, indicated in the top right corner of each panel, against the number of species in the plot. Lines represent linear model fits for pooled data (black) as well as each treatment (colours). Legend shows colour codes for treatments. Actual data points for each plot represented as grey dots.

The diversity-multifunctionality effect (DME) is sensitive to trophic complexity. In the pooled dataset, where all treatments were taken together, the DME increased and peaked at moderate thresholds, switching to negative effect at high thresholds, following predictions of the jack-of-all-trades effect (Fig 4). Trophic complexity had an effect on both the height and location of the peak DME (Fig 4). Comparing the curves of the four different treatments, we found that the addition of either the above-ground (INS) or litter (LIT) trophic level led to higher peak DMEs than the plant-only or full-complexity (BOTH, i.e., plants, litter, and above-ground fauna) communities, but all treatments peaked at similar intermediate thresholds (∼ 35-40%). The transition from positive to negative DMEs occurred between ∼ 40 – 60% thresholds for all trophic treatments except the full complexity treatment. Finally, the full trophic complexity treatment was the most distinct of the four treatments, having the lowest peak (at ∼10% threshold) with DME values that were consistently lower in magnitude. This treatment also showed the earliest switch to negative values, at roughly 20% function threshold, remaining largely negative across most threshold values.

**Figure 4.**
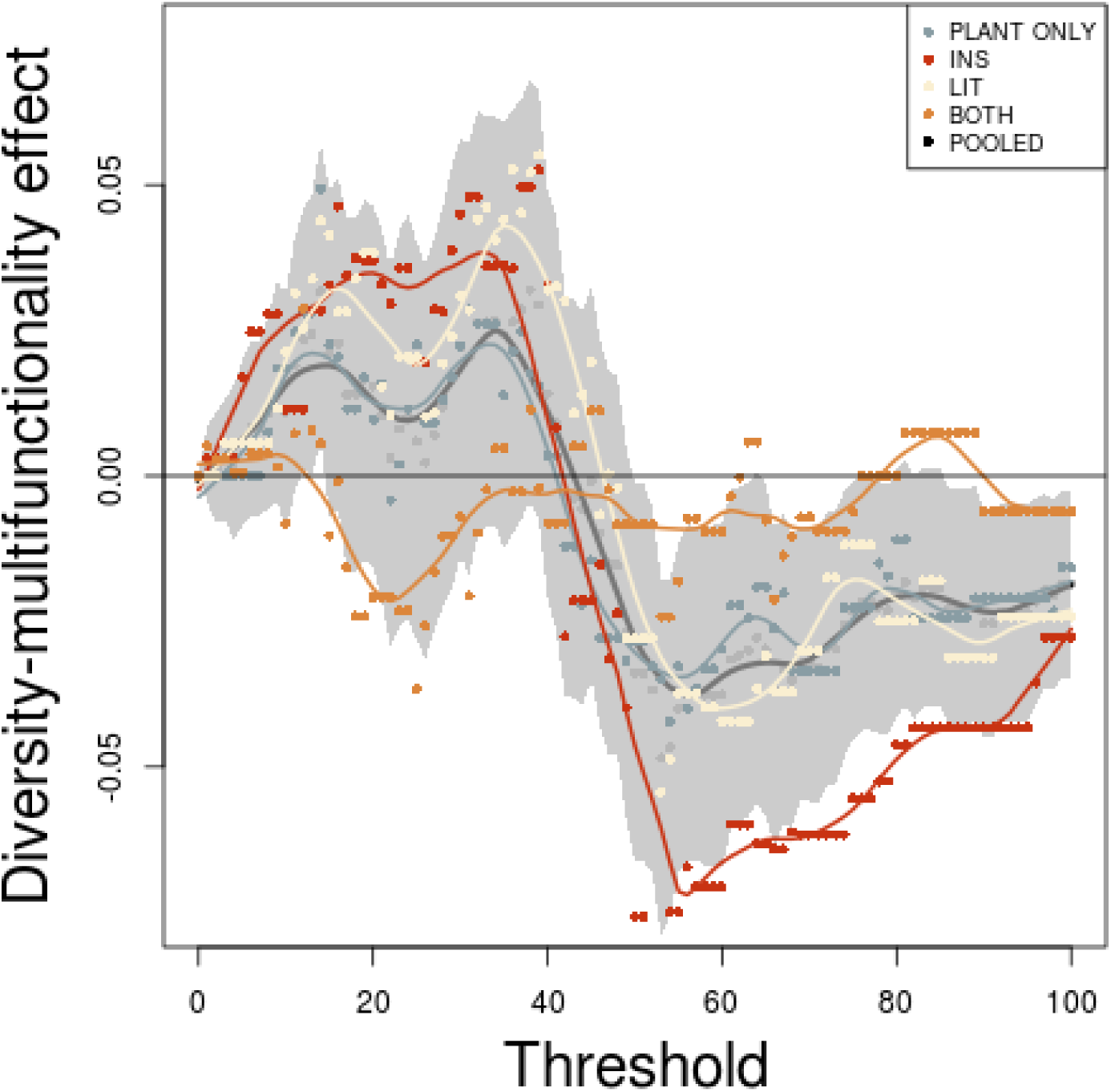
Effect of threshold on the biodiversity-multifunctionality effect (the DME). Each point represents the slope of the linear model between plant species richness and number of functions above the threshold. Each DME curve for each level of trophic complexity is plotted using a different colour, as presented in the key (top right), with the curve for the pooled dataset presented in black. The curves are smooth-spline interpolations. The grey polygon represents the bounds of the standard error of the slope in the pooled dataset.

## Discussion

In agreement with recent literature^7,8^, our results show that biodiversity is necessary for maintaining multiple functions from low to moderate function threshold values in grassland communities but at higher thresholds, increases in diversity are inversely associated with multifunctionality (i.e., the jack-of-all-trades effect) (Figs 3-4).

We advance understanding of biodiversity and multifunctionality, however, by also showing that this jack-of-all-trades and master-of-none pattern is sensitive to trophic complexity. Although the bottom-up effect of plant diversity on herbivore diversity, and potentially ecosystem functions, has been observed in long-term experimental data^17,19–21^, our study tests the top-down (aboveground fauna) and donor-controlled (litter fauna) effects of trophic complexity on multifunctionality. The simultaneous presence of aboveground and litter fauna trophic levels, the condition closest to natural systems where trophic complexity is common, showed a switch to negative diversity-multifunctionality effects (DMEs) at a very low threshold and maintained persistent negative DMEs throughout most thresholds values (Fig. 4). Any reduction in trophic complexity altered the diversity-multifunctionality relationship. Thus, both the location and height of peak DMEs are affected by trophic complexity, as suggested in our conceptual framework (Fig. 1).

The nature and direction of the trophic impact on DME depends on the trophic component considered. When examined across thresholds, both aboveground and litter fauna amplified the DME at low thresholds but at high thresholds, only aboveground fauna amplified the effect. This suggests that the presence of aboveground arthropods increases the contribution of biodiversity in providing ecosystem multifunctionality at moderate thresholds. This could be because plant diversity is known to decrease associational effects to herbivory damage^22, 23^ leading to an additional advantage of diversity in the presence of aboveground fauna.

In contrast to what we observed for multifunctionality, our analyses did not reveal impacts of trophic complexity on any single ecosystem function when aggregated across plant diversity treatments (Figs. 2-3). Further we did not observe any impacts of the trophic complexity treatments on average ecosystem function value at any level of plant diversity (Fig S1). Although the treatment with plants alone performed better at water retention along all thresholds (Fig. 2) and intermediate diversities (Fig S1), this observation alone cannot explain the differences between the treatments.

Despite the lack of significant effects on single ecosystem functions, we observe trophic complexity effects at the multifunctionality level. Our results also show that the presence of multiple trophic components decreases the magnitude of the DME across all thresholds, consistent with results from a long-term experiment from the same region demonstrating that non-producer trophic components obscure the diversity-productivity relationship due to a loss in complementarity effects with the addition of herbivores^17^. As the plants we used were taxonomically and functionally quite distinct, identity effects^24^, or unique contributions of individual species to the ecosystem functions considered, are also likely to be responsible for the nature of the biodiversity-multifunctionality relationship we observed.

Our findings have important implications for understanding the relationship between biodiversity and ecosystem functions. Plant diversity is currently understood to be critical to sustaining multifunctionality at, or below, moderate function threshold values, but our results show that such effects are influenced by trophic complexity. Given that declines in trophic complexity are widespread, occurring, for example, where agriculture leads to habitat simplification through the use of biocides and where trophic downgrading is occurring in terrestrial and marine habitats, most ecosystems are experiencing declines in both producer diversity and trophic complexity. Sustaining a broad spectrum of ecosystem functions and the services they provide will require more plant diversity in the face of widespread trends in trophic simplification.

## Methods

### Experimental Methods

We used data from a year-long experiment on grassland mesocosms in a tall-grass prairie, part of the Cedar Creek Ecosystem Science Reserve, Minnesota. The experimental design was factorial with 100 pots, 1m in diameter, grown inside netted insect exclosures. Each pot was maintained at one of 5 levels of plant diversity: 1, 2, 4, 8 or 16 species. The species used in this experiment (Table S1) were native perennial species used in other experimental studies from the site^17,25,26^. Pots with incomplete data on species identity were excluded from this analysis, resulting in a sample size of 94. The plant diversity treatments were crossed with trophic complexity treatments: plants and aboveground mesofauna (primarily insects and other invertebrates, INS), plants and litter mesofauna (LIT), plants and both aboveground and litter mesofauna (BOTH), or plants only (i.e., no aboveground or litter fauna, NONE). These treatments were achieved by first applying a pesticide treatment on all the pots and removing all aboveground and litter fauna. 25% of the pots in each treatment were allowed to be recolonised by aboveground fauna only, 25% by liter fauna only, 25% by both aboveground and litter fauna, and 25% were plants only, leading to the four community complexity treatments. The experiment was run for a year from July 2000 to July 2001 when final biomass and soil water retention were measured and biomass recovered in each pot after harvest was measured a year later.

### Ecosystem function measurements

Four candidate ecosystem functions were analysed: (i) Aboveground biomass: the total aboveground biomass in each pot was measured in 2001, after one year of experiment. (ii) Belowground biomass: the total belowground biomass in each pot was measured after one year of experiment (iii) Water retention: the time taken for a fixed volume of water to flow into a collection flask at the bottom of the pot was measured at the end of one year of experiment (iv) Biomass recovery: the biomass recovered one year post harvest was measured to quantify biotic control over ecosystem resilience. Recovery was calculated as the ratio of the total recovered biomass to the biomass prior to disturbance (aboveground and belowground biomass). A principal component analysis of functions to test correlations resulted in water retention close to PC1 and aboveground biomass close to PC2 and the vector of biomass post-harvest was in a similar direction as the biomass in the previous year.

For each function, the decay curves for each trophic treatment was plotted along a standardised range of threshold values. These were visually examined for differences.

### Multifunctionality

To assess multifunctionality across diversity treatments, measurements of four ecosystem functions – aboveground biomass, belowground biomass, water retention and biomass recovery after harvest - were chosen and analysed using published methodology of the threshold approach^18^. To this end, the maximum value for each ecosystem function was calculated as the mean of the five highest function values in the entire experiment. Each ecosystem function in a pot was then standardised between this maximum and the minimum value in the experiment. For every 5% between 0 and 100, each pot was scored for the number of functions maintained above that threshold. For each threshold, the slope of linear model between the number of functions maintained above threshold and the manipulated plant diversity in the community was defined as the diversity effect on multifunctionality (DME). DME was analysed for the pooled dataset as well as the dataset split into the four trophic complexity treatments. The magnitude of the peak and the point at which the curve of biodiversity effect vs. threshold crosses the x-axis for each of the treatments were examined in comparison with the pooled data for reasons described in the introduction.

