## Supplemental Table 1 and Figure S1 for "Trophic complexity alters the diversity-multifunctionality relationship in experimental grassland mesocosms"

Supplementary information

Table S1. List of plant species used for the experiment

| Sl No | Species |
| --- | --- |
| 1 | *Achillea millefolium* |
| 2 | *Andropogon gramine* |
| 3 | *Anemone cylindrica* |
| 4 | *Asclepias tuberosa* |
| 5 | *Buchloe dactyloides* |
| 6 | *Chondrosum gracile* |
| 7 | *Coreopsis palmata* |
| 8 | *Elymus canadensis* |
| 9 | *Euphorbia corollata* |
| 10 | *Koeleria cristata* |
| 11 | *Liatris aspera* |
| 12 | *Panicum virgatum* |
| 13 | *Rudbeckia hirta* |
| 14 | *Schizachyrium scoparium* |
| 15 | *Solidago nemoralis* |
| 16 | *Sorghastrum nutans* |
| 17 | *Sporobolus cryptandrus* |
| 18 | *Symphyotricum subulatum* |

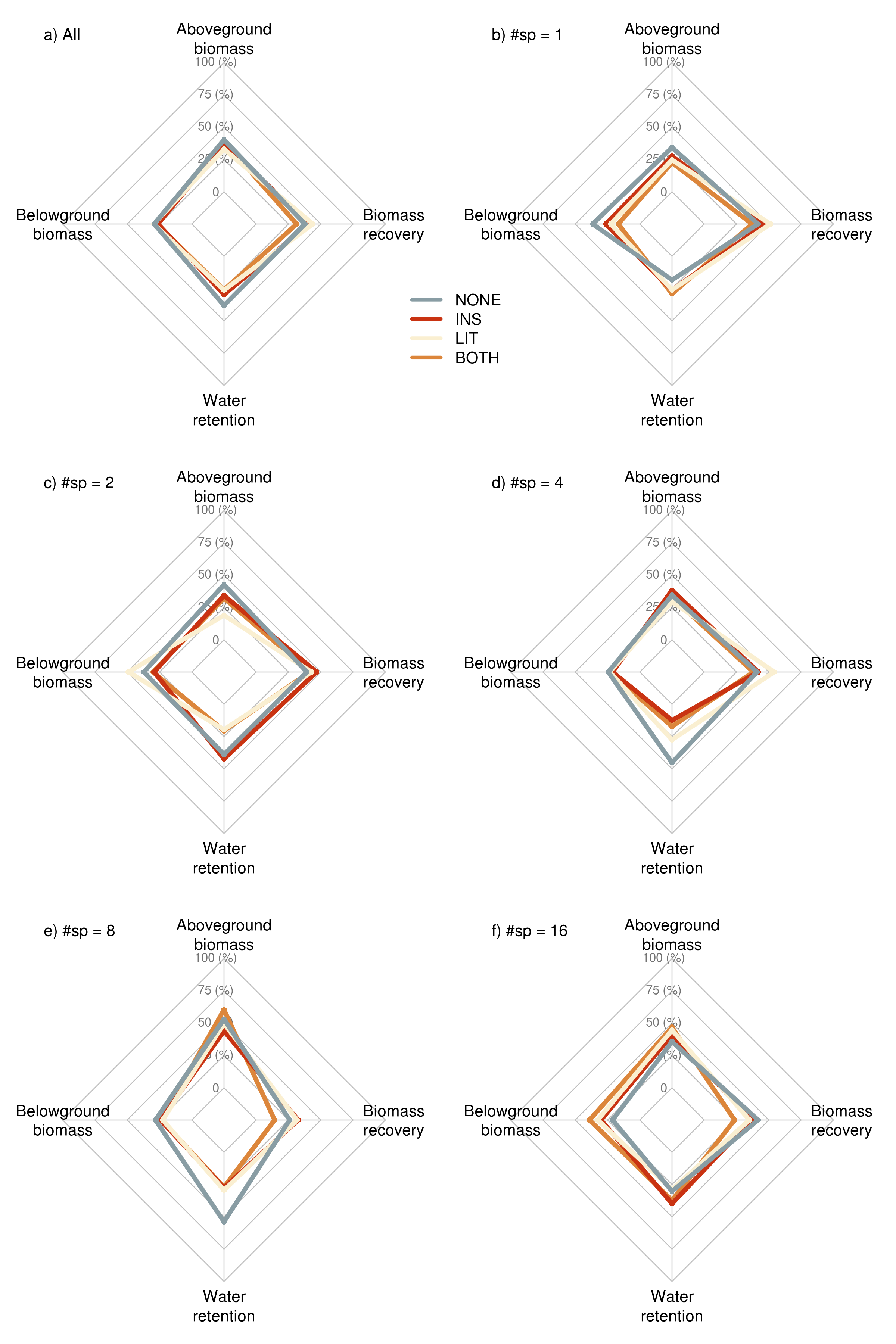

Figure S1 : Spider plot showing the average percent of each function maintained by communities with different number of species and trophic complexity. a) The average percent of each function maintained by all communities in the experimental setup. The next panels represent average percent of each function maintained by communities with b) 1 plant species c) 2 plant species d) 4 plant species e) 8 plant species and f) 16 plant species. The four colours represent the four trophic treatments (PLANT ONLY, INS, LIT and BOTH) within each plant diversity treatment. The points along each axis is calculated as the mean of each function scaled between the maximum and minimum value in the entire experiment.
